# Human Papillomavirus Genome Copy Number Is Maintained by S Phase Amplification, Leakage to the Cytosol during Mitosis and Lysosomal Degradation in G1 Phase

**DOI:** 10.1101/2022.06.24.497574

**Authors:** Malgorzata Bienkowska-Haba, Katarzyna Zwolinska, Timothy Keiffer, Martin Sapp

## Abstract

The current model for human papillomavirus (HPV) replication is comprised of three modes of replication. Following infectious delivery, the viral genome is amplified during the establishment phase to reach up to some hundred copies per cell. HPV genome copy number remains constant during the maintenance stage. Differentiation of infected cells induces HPV genome amplification. Using highly sensitive *in situ* hybridization (DNAscope) and freshly HPV16-infected as well as established HPV16-positive cell lines, we observed that viral genome is amplified in each S phase of undifferentiated keratinocytes cultured as monolayers. Nuclear viral genome copy number is reset to pre-S phase levels during mitosis. The majority of viral genome fails to tether to host chromosomes and is lost to the cytosol. Cytosolic viral genome gradually decreases during cell cycle progression. Loss of cytosolic genome is blocked in presence of NH_4_Cl or other drugs interfering with lysosomal acidification, suggesting the involvement of autophagy in viral genome degradation. These observations were also made with HPV31 cell lines obtained from patient samples. Cytosolic viral genome was not detected in UMSCC47 cells carrying integrated HPV16 DNA. Analyses of organotypic raft cultures derived from keratinocytes harboring episomal HPV16 revealed the presence of cytosolic viral genome as well. We conclude that HPV maintains viral genome copy number by balancing viral genome amplification during S phase with loss of viral genome that is lost to the cytosol during mitosis. It seems plausible that restrictions to viral genome tethering to mitotic chromosomes resets genome copy number in each cell cycle.

**IMPORTANCE:** HPV genome maintenance is currently being thought to be achieved by regulating expression and activity of viral replication factors E1 and E2. In addition, the E8^E2 repressor has been shown to be important for restricting genome copy number by competing with E1 and E2 for binding to the viral origin of replication and by recruitment of repressor complexes. Herein, we demonstrate that HPV viral genome is amplified in each S phase. Nuclear genome copy number is reset during mitosis by a failure of the majority of genomes to tether to mitotic chromosomes. Rather, they accumulate in the cytoplasm of freshly divided cells. Cytosolic viral DNA is quickly degraded in G1 in a lysosome dependent manner contributing to the genome copy reset. Our data imply that the mode of replication during establishment and maintenance is the same and further suggest that restrictions to genome tethering significantly contributes to viral genome maintenance.

## INTRODUCTION

High-risk human papillomaviruses (HPV) such as HPV16 are causative agents of squamous cell carcinoma of the oropharynx and anogenital tract, including the cervix. They establish infection in the basal cell compartment of the stratified epithelia of the skin and mucosa by delivering viral genome to the nucleus of target cells. Often, HPV infections are subclinical with no overt clinical symptoms; the majority of HPV-induced clinical disease can be categorized as benign self-limited outgrowth of the skin and mucosa presenting as papilloma or warts. Cellular transformation events are often associated with HPV genome integration and require infection by the high-risk HPV types most commonly of species 7 and 9 of the alpha genus (1).

The viral genome of approximately 8,000 bp is retained in the infected cell as an extrachromosomal double stranded DNA. This viral genome is circular and associated with histones. Based on the observation that cells harbor up to some hundred extra-chromosomal copies, it is assumed that the viral genome is amplified during infection establishment. During latency and genome maintenance of keratinocytes in the basal cell compartment, viral genome copy number is maintained. However, when keratinocytes enter the terminal differentiation process, viral genome is amplified followed by expression of capsid proteins (1, 2).

HPV encodes two proteins essential for viral genome replication. The E1 protein binds to the origin of replication, directed by the E2 protein. E1 binding results in melting of the double stranded DNA and allows for recruitment and binding of cellular replication factors to initiate replication. E1 protein also functions as a helicase during the elongation phase, whereas E2 protein dissociates from the replication complex (3, 4). In addition, regulatory viral factors such as the E8^E2 repressor have been shown to regulate genome copy number (5, 6). During maintenance, genome is replicated via the bi-directional mode (1, 7, 8). Several reports have suggested that differentiation-induced genome amplification utilizes a rolling circle mechanism based mainly on data derived from the analysis of two-dimensional gel electrophoresis (8-11). However, cellular factors driving this mode of replication have not been positively identified. Little is known about the mode of replication during the establishment phase.

Genome copy number is thought to be regulated by: viral repressors such as E8^E2, both at the transcriptional and posttranslational level (5, 12); regulation of expression levels of the viral replication factors E1 and E2 (1, 7); and post-translational modification of the E1 protein such as proteolytic cleavage and phosphorylation (3). However, these findings were mostly based on bulk cell analyses using quantitative PCR and Southern blot. The lack of methodology allowing detection of low copy number viral genome on a single cell basis prevented a more detailed analysis on a single cell level. Herein, we use highly sensitive fluorescent *in situ* hybridization to follow the fate of viral genome over the cell cycle of both freshly infected and immortalized keratinocytes. We observed that viral genome maintenance is achieved by unlicensed genome amplification during S phase, followed by loss to the cytosol during mitosis, and subsequent lysosomal degradation in G1.

## RESULTS

### Genome copy number is highly heterogeneous in established and freshly infected keratinocytes

When we stained immortalized HPV16 harboring keratinocytes at 7- and 45-days post infection (dpi) for detection of viral DNA using a highly sensitive fluorescent *in situ* hybridization technique, DNAscope, and an L1-specific probe, we noticed that the number of fluorescent puncta representing viral genome was highly variable. Surprisingly, some cytoplasmic puncta were detected in addition to nuclear puncta, mostly in cells harboring high genome copy numbers (Fig 1B, 1E). Quantification of the cytoplasmic and nuclear puncta revealed that approximately 35% of detected genome signals localized to the cytoplasm of interphase cells; In mitotic cells, around 70% of genome puncta were not associated with mitotic chromosomes (Fig 1C, 1F). No specific HPV16 genome puncta were detected in uninfected control primary keratinocytes suggesting that the probe was specific for HPV16 (Fig 1A, 1C). Although we see similar results for freshly infected (7 days post-infection) and long-infected (>45dpi) cells, not all cells for the 7dpi condition were infected and thus the overall number and size of HPV genome puncta for these freshly-infected cells were somewhat lower (Fig 1B, 1E). To control for the completely unexpected finding of cytoplasmic viral DNA, we confirmed that RNAse treatment completely degraded RNA as shown here for the cellular household gene cyclopihlin B (CyPB) (Fig 2A). We also confirmed that our protocol utilized for DNAscope does not allow detection of RNA after RNAse treatment (Fig 2B) and also that DNase I treatment prior to hybridization results in complete loss of viral genome detection in both nucleus and cytoplasm (Fig 2C). We then tested our protocol using the UMSCC47 cell line, which has the HPV16 viral genome integrated into the host genome (13), and we found that detection of viral genome was restricted to the nucleus (Fig 2D). RNAscope of UMSCC47 cells shows the expected pattern of staining. Since we are using HPV31-harboring cell lines in subsequent experiments, we also tested specificity of probes and found no cross-detection when using our HPV16 and HPV31 specific probes (Fig 2E).

**FIG 1.**
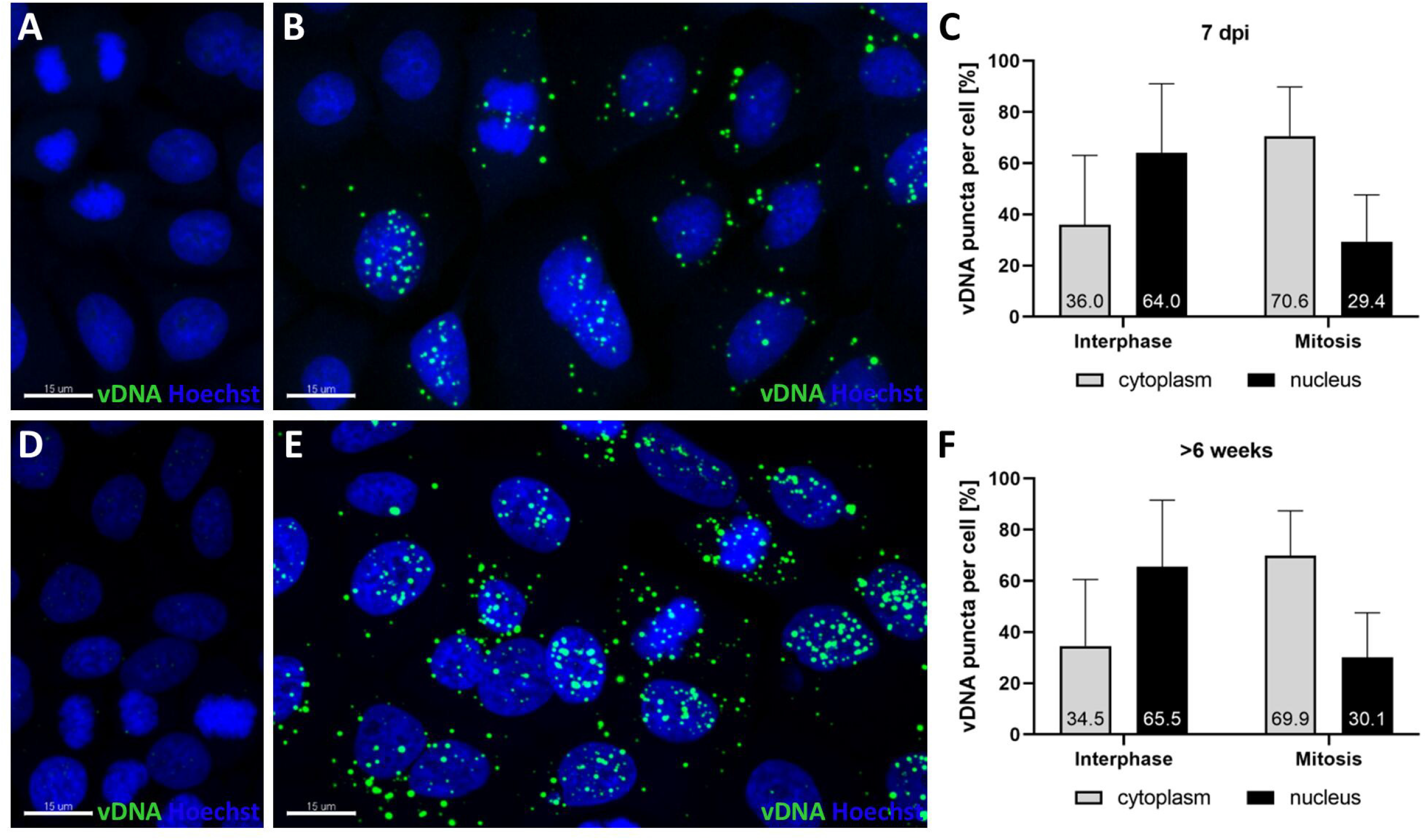
Cellular localization of HPV16 genome. Viral genomes detected by highly sensitive *in situ* DNA hybridization (DNAscope) with the HPV16 L1 probe in human foreskin keratinocytes infected with WT HPV16 and cultured in monolayer for 7 dpi (B) and above 6 weeks post infection (E), images of uninfected control HFKs alongside the 7dpi HFKs (A) and >6weeks post-infected HFKs (D) are also included. Average percentage of viral genomes present in nuclei and cytoplasm of mitotic and interphase cells 7 dpi (C) or in long-term culture (F). Bars represent standard deviation of counts taken with 371 interphase and 66 mitotic cells coming from three independent experiments conducted at day 7 (C), and 474 interphase and 67 mitotic cells at >6 weeks post-infection (F) from three independent experiments. Control uninfected keratinocytes were subjected to DNAscope with HPV16 probe. All pictures are presented as maximum projection of at least 12 z-stacks; scale bars 15 μm.

**FIG 2.**
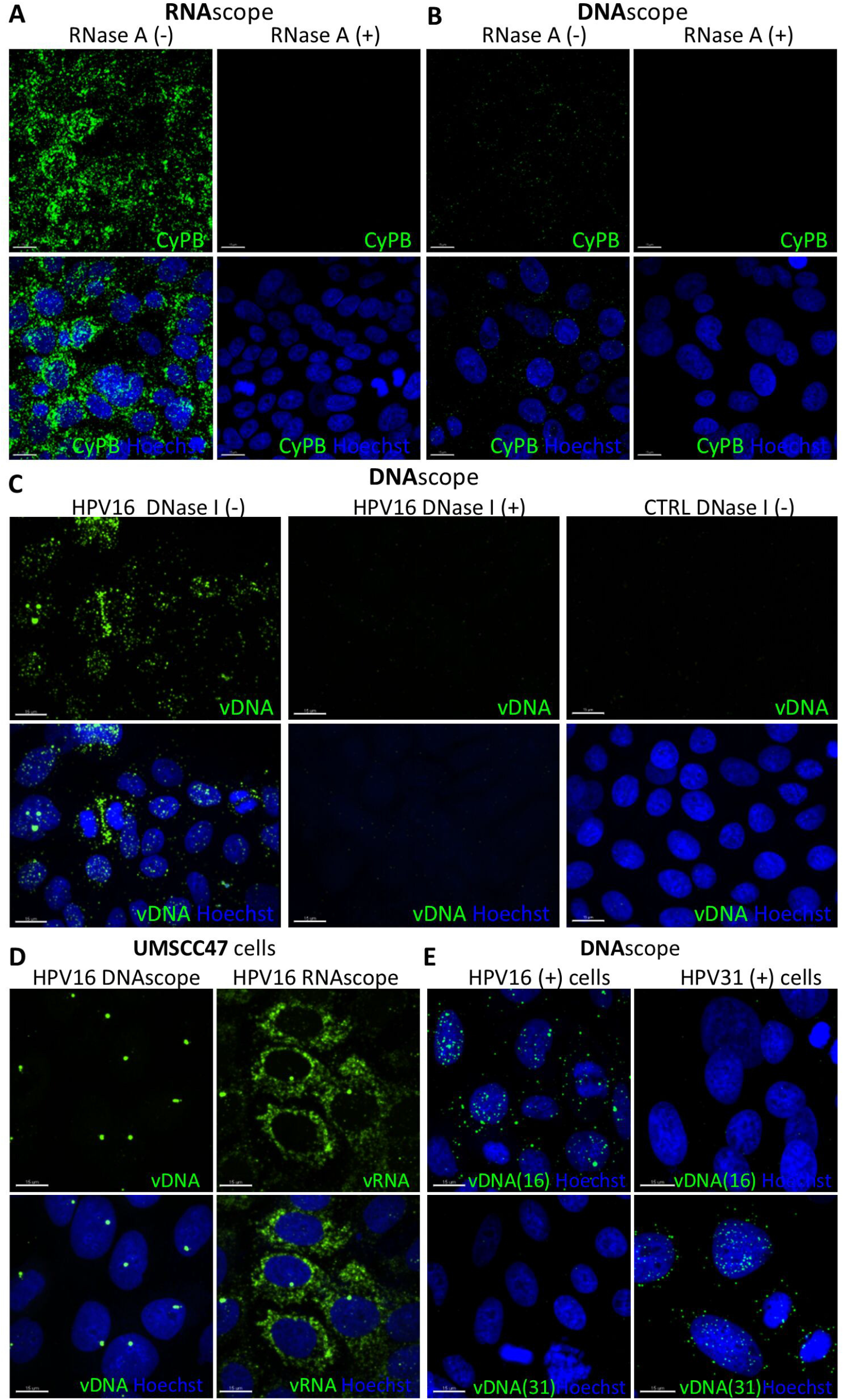
Controls for specificity of highly sensitive *in situ* hybridization of RNA (RNAscope) and DNA (DNAscope). Uninfected human foreskin keratinocytes were subjected to hybridization with the probe for human cyclophilin B (CyPB) in RNAscope (A) and DNAscope conditions (B), in the presence or absence of RNase A. (C) DNAscope of HPV16 infected cells treated with DNase I compared to untreated infected or uninfected cells. (D) DNAscope with HPV16 L1 and RNAscope with E6/E7 probes in UMSCC47 cells containing integrated HPV16 genomes. (E) HPV16-or 31-positive cells subjected to DNA scope with HPV16 and 31-specific probes. All pictures are presented as maximum projection of at least 12 z-stacks; scale bars 15 μm.

### Localization of viral genome is dependent on the cell cycle phase

When we focused our analysis on mitotic cells, we observed that only a small subset of viral genome segregated with the mitotic chromosomes and a large fraction of viral genome dissociated from cellular chromosomes, often lining up around microtubules in telophase (Fig 3). While most viral genome still colocalized with host cell chromosomes during prophase, separation from mitotic chromosomes became first evident in metaphase and continued to increase through anaphase. This separation from mitotic chromosomes seen with HPV genomes was most prominent during telophase and cytokinesis.

**FIG 3.**
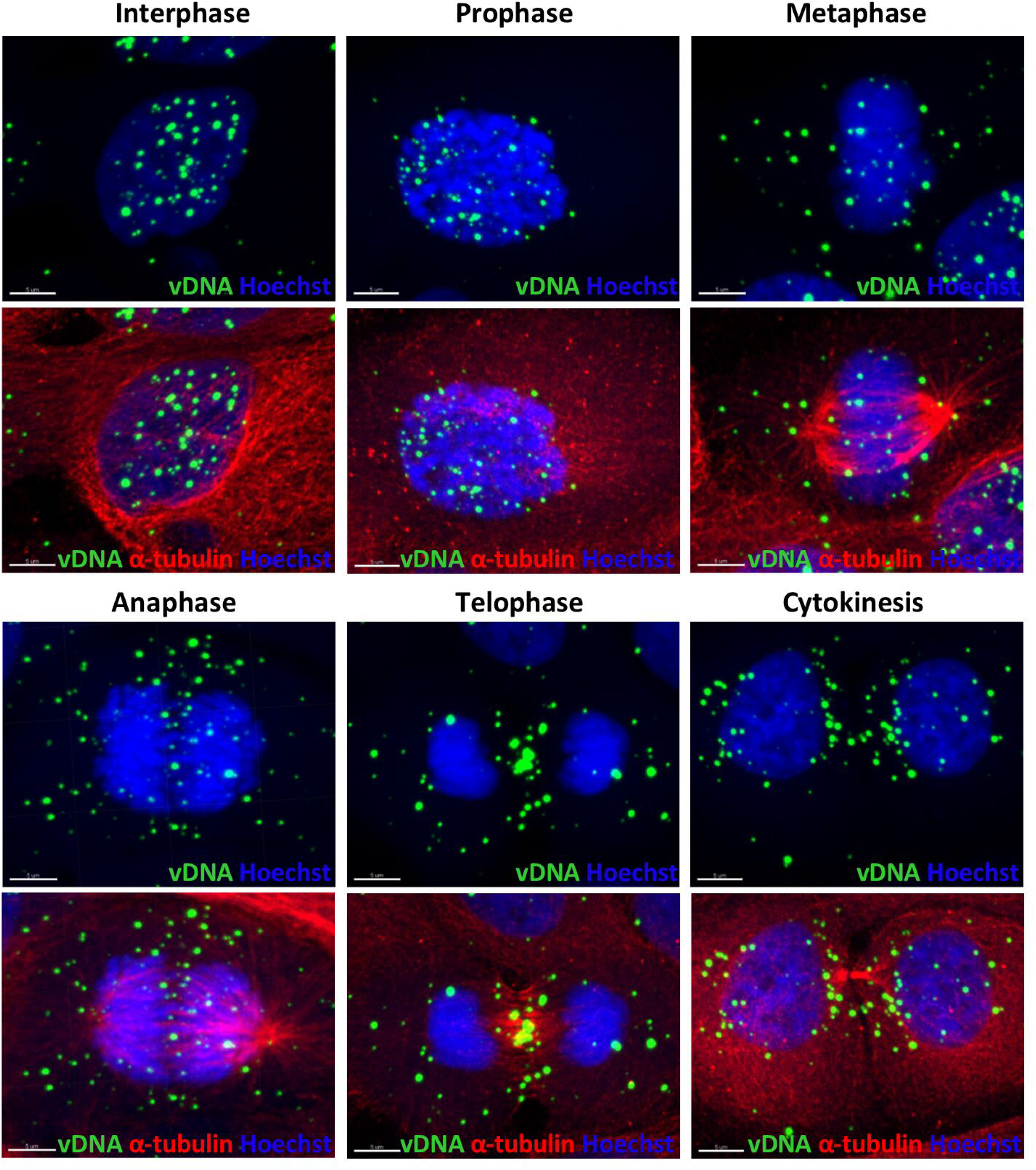
Representative image of intracellular distribution of HPV16 genomes in different phases of mitosis and an example of interphase cell. All pictures are presented as maximum projection of at least 12 z-stacks, scale bars 5 μm. Prophase cells show mostly nuclear localization of viral puncta, however, in metaphase part of viral genomes become separated from mitotic chromosomes. Cytoplasmic localization of viral DNA is even more pronounced in later phases, especially telophase and cytokinesis.

Very often, the number of virus-specific foci were higher in mitotic cells compared to non-dividing cells. Therefore, we hypothesized that viral genome is amplified during S and possibly G2 phase and that genome localizing to the cytosol is degraded in G1. To test this, we combined DNAscope with EdU pulse labeling to label cells in S phase along with immunofluorescent staining for cyclin B1. To distinguish between early and late G1 stages, we counted the number of nucleoli, which decrease as the interphase cells mature. This allowed us to identify cells in early G1 (small, numerous nucleoli), mid to late G1 (up to 3 nucleoli), early S phase (positive for EdU incorporation, negative for cyclin B1), mid and late S phase (positive for EdU and increasing cyclin B1 signal strength), G2 (positive for cyclin B1) and mitotic cells based on mitotic figures. Figures 4A and B depict a representative image of HPV16-infected keratinocytes at 45 dpi and an uninfected control, respectively. In Figure 4C, we displayed representative cells during the respective cell cycle stages. We quantified the total number of virus-specific foci residing in the nucleus and the cytosol for each cell cycle stage at day 7 (Fig 5A) and >day 45 (Fig 5B) post-infection of primary keratinocytes with HPV16. We observed a decrease of cytosolic puncta from early G1 to late G1. In contrast, nuclear viral genome puncta number per cell increased during S phase and reached a peak in G2 phase. During this time, not only the number of puncta, but also size and intensity of nuclear puncta, increased. Even though the overall number of viral genome puncta is higher at >45 dpi, there is no qualitative differences at 7 and >45 dpi. The increase is restricted to puncta residing in the nucleus, suggesting genome amplification. Indeed, some EdU incorporation is observed in replication foci in S and even in G2 phase (Fig 5C), confirming reports by others that viral genome replicates throughout G2 (14). In mitosis, only a small fraction of viral genome remained associated with mitotic chromosomes.

**FIG 4.**
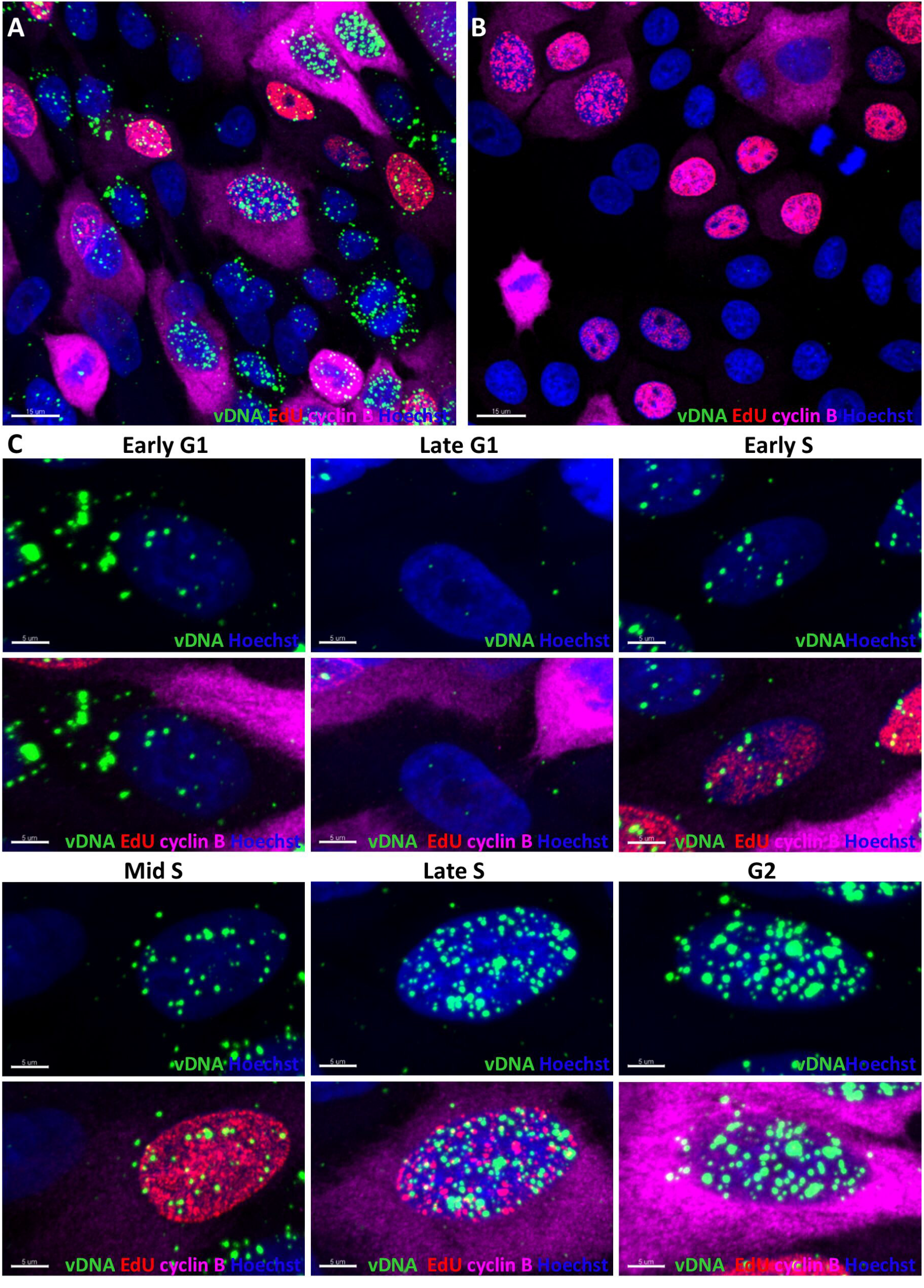
Nuclear and cytoplasmic HPV16 genome distribution during interphase of cell cycle of infected primary human keratinocytes. Episomal HPV16 harboring keratinocytes (A) and uninfected controls (B) were EdU labeled for 30 min and stained for HPV16 genome by DNAscope(green), EdU by ClickIt reaction (red) as an indicator of S-phase, and cyclin B by indirect immunofluorescence (magenta). Nuclei were stained by Hoechst. Cyclin B positive but EdU negative cells were recognized as G2 or late S; G1 cells were negative for both, EdU and cyclin B. Early and late G1 stages were distinguished based on the presence, number and proportions of nucleoli (small, numerous nucleoli in early G1 and up to 3 big nucleoli in late G1). Panel C depicts examples for each cell cycle phase. All pictures are presented as maximum projection of at least 12 z-stacks; scale bars 15 μm (A, B) and 5 μm (C).

**FIG 5.**
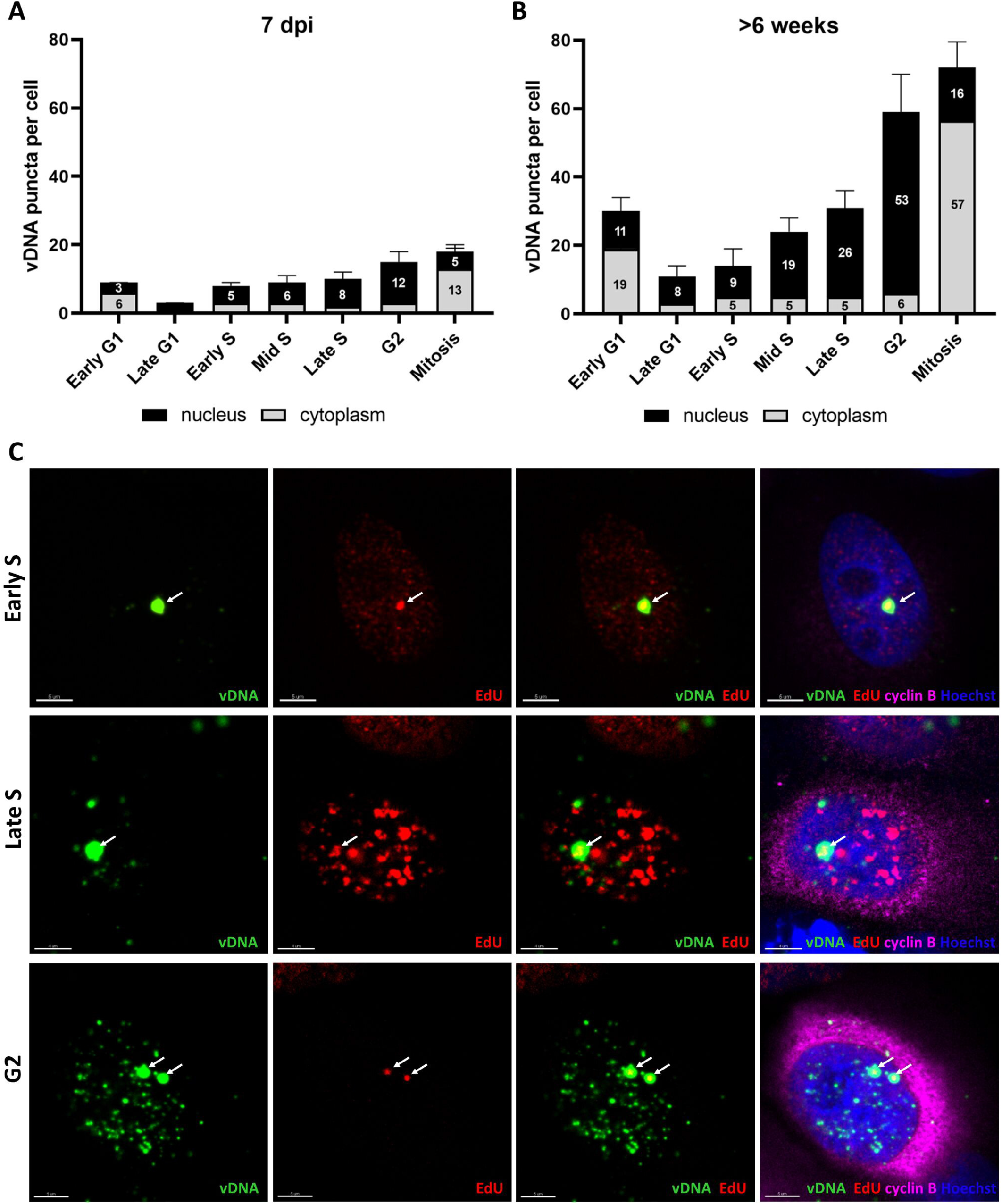
Quantification of intracellular HPV16 genome distribution in freshly infected and immortalized HFK. HFK were stained as described in Fig 4 at 7 dpi (A) and above 6 weeks after infection (B) and nuclear and cytoplasmic viral genome foci were quantified as a function of the cell cycle phase. Viral genome amplification is reflected by increasing number of HPV16 genome puncta in nuclei of cells in S/G2 phases. The number of viral DNA puncta in the cytoplasm is highest in M and early G1 phase and gradually decreases during cell phase progression. The data are presented as median puncta number per cell with 95% confidence interval. At least 60 cells per cell cycle phase from three independent experiments were included in the analysis. (C) Viral amplification foci colocalizing with EdU staining reflects active replication during early S (upper panel), late S (middle panel) and G2 stage (lower panel) of the cell cycle. Images are single slices chosen from at least 12-slices from z-stacks, scale bars 5 μm.

### Cytosolic viral genome is also observed in long-term immortalized keratinocytes established from low-grade lesions or after genome transfection and selection

To address whether our observation is specific for infected keratinocytes and HPV16, we extended our analysis to immortalized HPV16 and HPV31 cell lines obtained after transfection of viral genome and outgrowth and to the CIN612-9E cell line, which was obtained from a cervical lesion and harbors episomal HPV31 (Fig 6). We observed a similar distribution of viral genomes in all analyzed cell lines, suggesting that the presence of cytosolic HPV DNA is not due to the different mode of viral genome delivery or HPV type-dependent (Fig 6B). Quantification of viral genome specific nuclear and cytoplasmic puncta is shown in Fig 6D-F. To ascertain that the HPV31 probe allowed specific detection of HPV31, we also performed RNAscope for all samples (Fig 6A) and used uninfected primary keratinocytes as control (Fig 6C).

**FIG 6.**
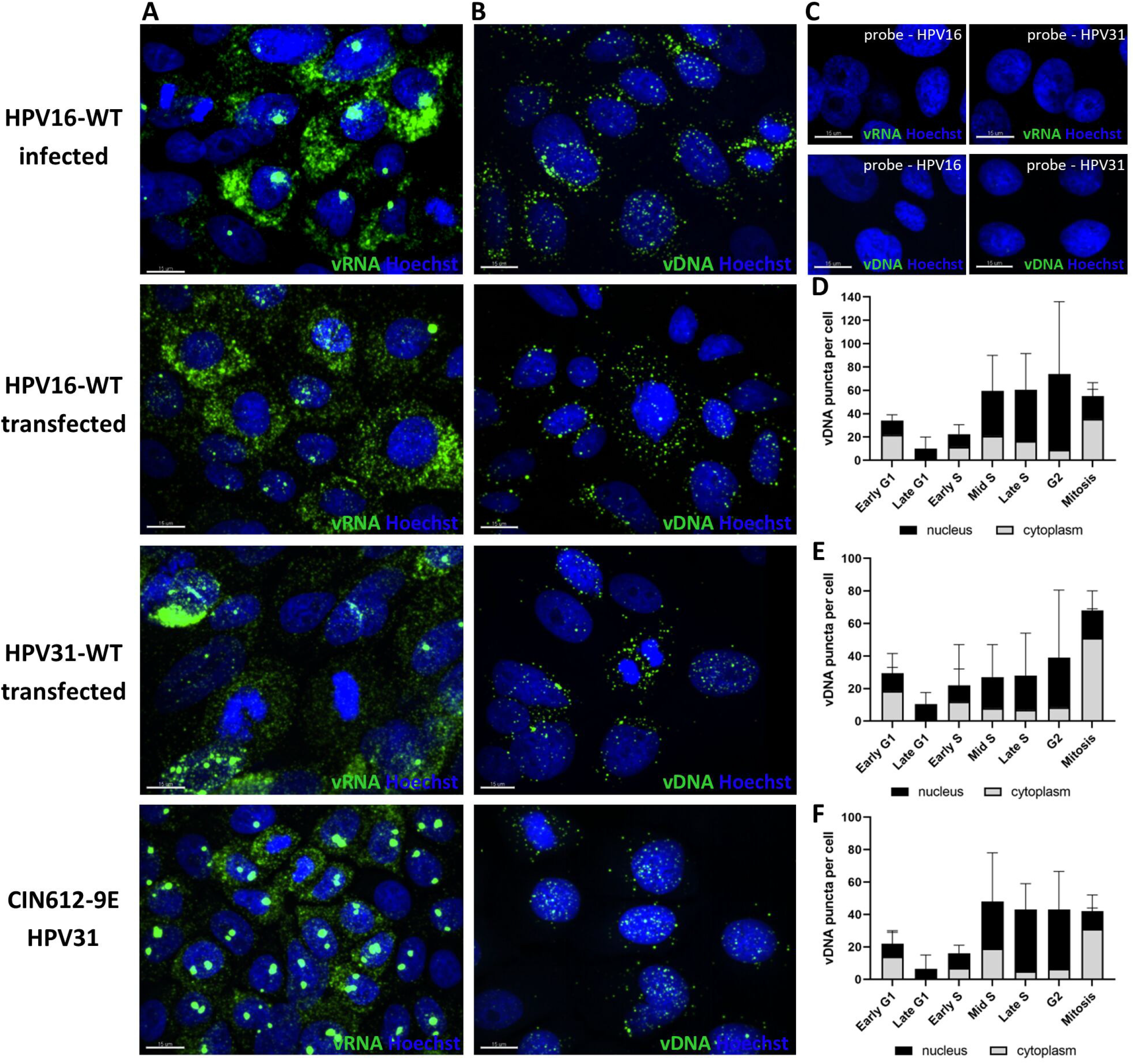
Common pattern of viral genome distribution and amplification in HPV16- and 31-positive keratinocytes. Immortalized HPV16-positive cells obtained after infection (HPV16-WT infected) or viral genome transfection of HFK (HPV16-WT transfected) as well as HPV31-positive cells derived from viral genome transfection (HPV31-transfected) or from a natural lesion (CIN12-9E line) were stained and analyzed as described in Fig 4 and 5. Panel B shows viral DNA detection with highly sensitive *in situ* hybridization DNAscope with type-specific HPV16-L1 and HPV31-E6/E7 probes. Viral RNA detection with HPV probes covering HPV16 and 31 E6/E7 sequences (panel A) is shown as a comparison of RNA detection patterns. (C) Control uninfected cells subjected to RNAscope and DNAscope with HPV16 or 31 specific probes. All pictures are presented as maximum projection of at least 12 z-stacks; scale bars 15 μm. (D-F) Viral genome quantification as function of the cell cycle in keratinocytes derived from transfection with HPV16 (D) or HPV31 (E) or CIN12-9E cell line (F).

To test whether the observation of loss of nuclear viral genome to the cytosol during mitosis is specific for human foreskin keratinocytes (HFKs) cultured in monolayer or is also observed in 3D cultures, we analyzed raft cultures grown from HPV16-infected keratinocytes (Fig 7A and B). We combined DNAscope with immunofluorescent staining for the S phase marker MCM7. Control organotypic rafts grown from uninfected primary keratinocytes confirmed specificity of the HPV16-specific DNAscope (Fig 7C and D). We found approximately 31% and 45 % of HPV16 puncta to reside in the cytoplasm of interphase and mitotic cells, respectively (Fig 7E).

**FIG 7.**
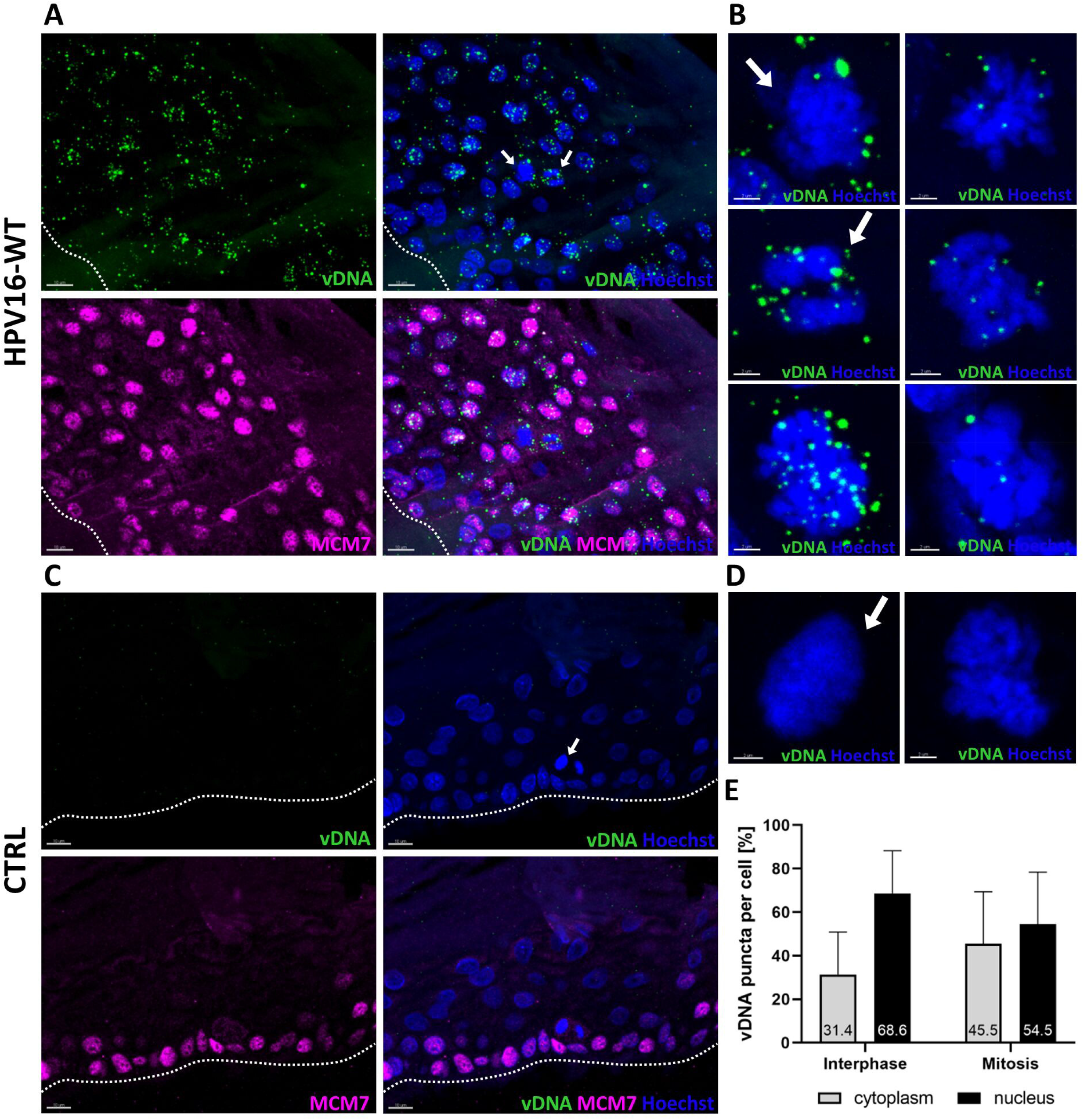
Intracellular localization of HPV16 genome in organotypic raft cultures. (A) DNAscope with HPV16 L1 probe of HFK infected with HPV16 grown in organotypic rafts shows both cytoplasmic and nuclear localization of viral genomes. (B) Examples of mitotic figures in infected rafts. (C,D) Control of uninfected cells subjected to DNAscope (C) and select mitotic figures (D). Images are presented as maximum projections from at least 12-slices z-stacks; scale bars 10 μm (A, C) and scale bars 2 μm (B, D). White dot line underline raft basal layer. (E) Quantification of viral genomes localizing to nuclei and cytoplasm of mitotic or interphase cells. Bars represent standard deviation of measurements taken of 70 interphase and 34 mitotic cells.

### Degradation of cytosolic viral genome can be blocked by inhibitors of lysosomal acidification

The decrease of cytosolic viral genome puncta during G1 suggested that the viral genome is degraded. To test a possible involvement of lysosomes, we treated HPV16-harboring HFK with ammonium chloride (NH_4_Cl), a potent inhibitor of lysosomal acidification. After 16 hours, cells were pulse labeled with EdU and processed for HPV16-specific DNA scope combined with cyclin B-specific immunofluorescence (Fig 8A). We observed a significant increase in cytoplasmic foci in treated cells as compared to untreated control at all cell cycle phases except for mitosis (Fig 8B). Similar results were obtained using Bafilomycin A1 and chloroquine as inhibitors of lysosomal acidification (Fig 9).

**FIG 8.**
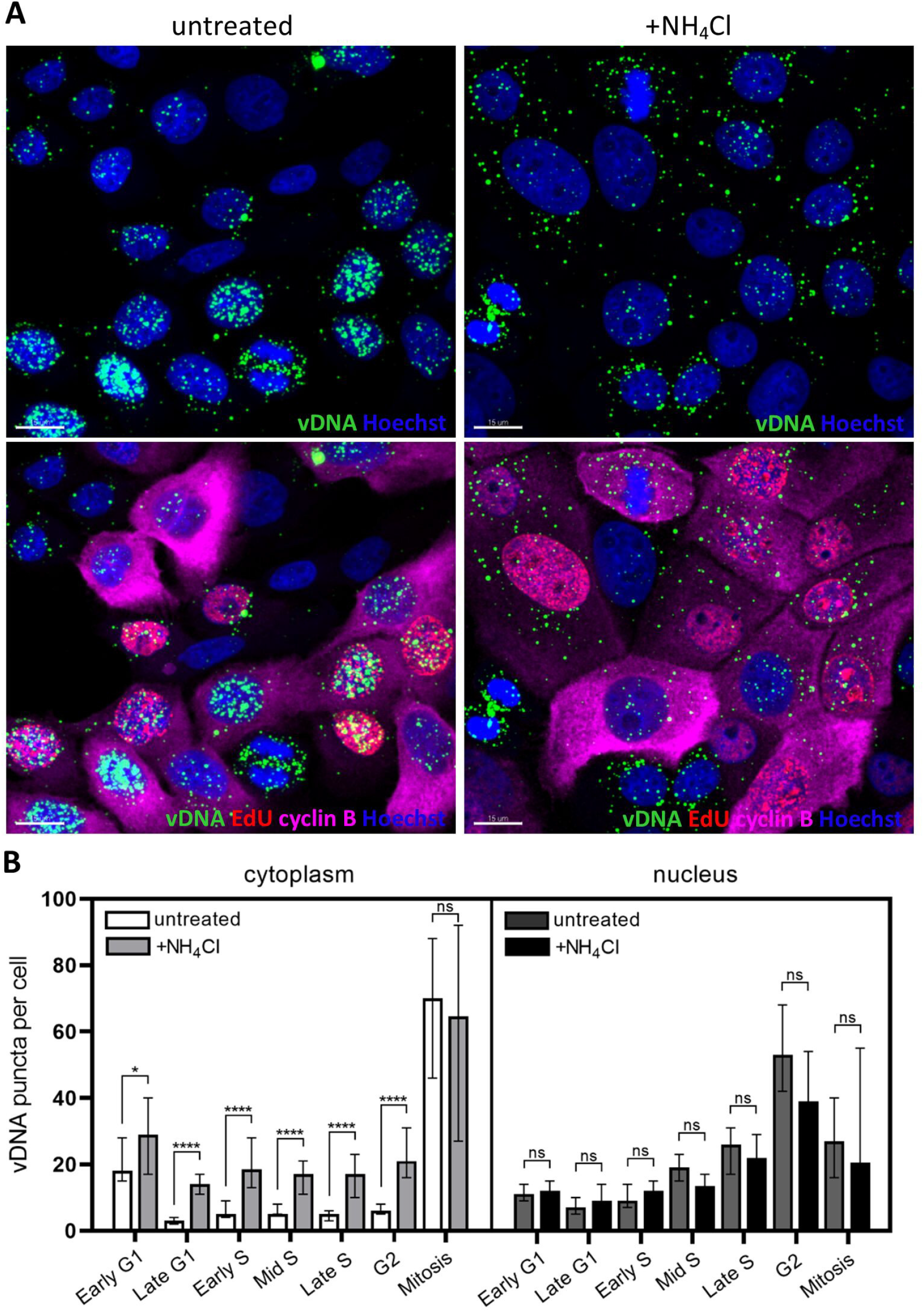
Ammonium chloride treatment blocks HPV genome degradation. HPV16 genomes detected in nuclei and cytoplasm of infected long-term cultured keratinocytes (>6 weeks) untreated or treated for 16-18 hours with lysosome pH-raising agent ammonium chloride (20 mM). Cells were stained and viral genome puncta were quantified as described in Fig 4 and 5. (A) Representative image of infected treated or untreated cells. Images are present as maximum projection from at least 12-slices z-stacks; scale bars 15 μm. (B) Quantification of cytoplasmic and nuclear HPV16 genome puncta in NH_4_Cl-treated and untreated cells suggesting involvement of lysosomes in degradation of viral DNA in cytoplasm. Data are presented as median viral puncta with 95% confidence interval and come from analysis of at least 60 cells per cell cycle phase in interphase and over 35 cells per mitosis from 3 independent experiments.

**FIG 9.**
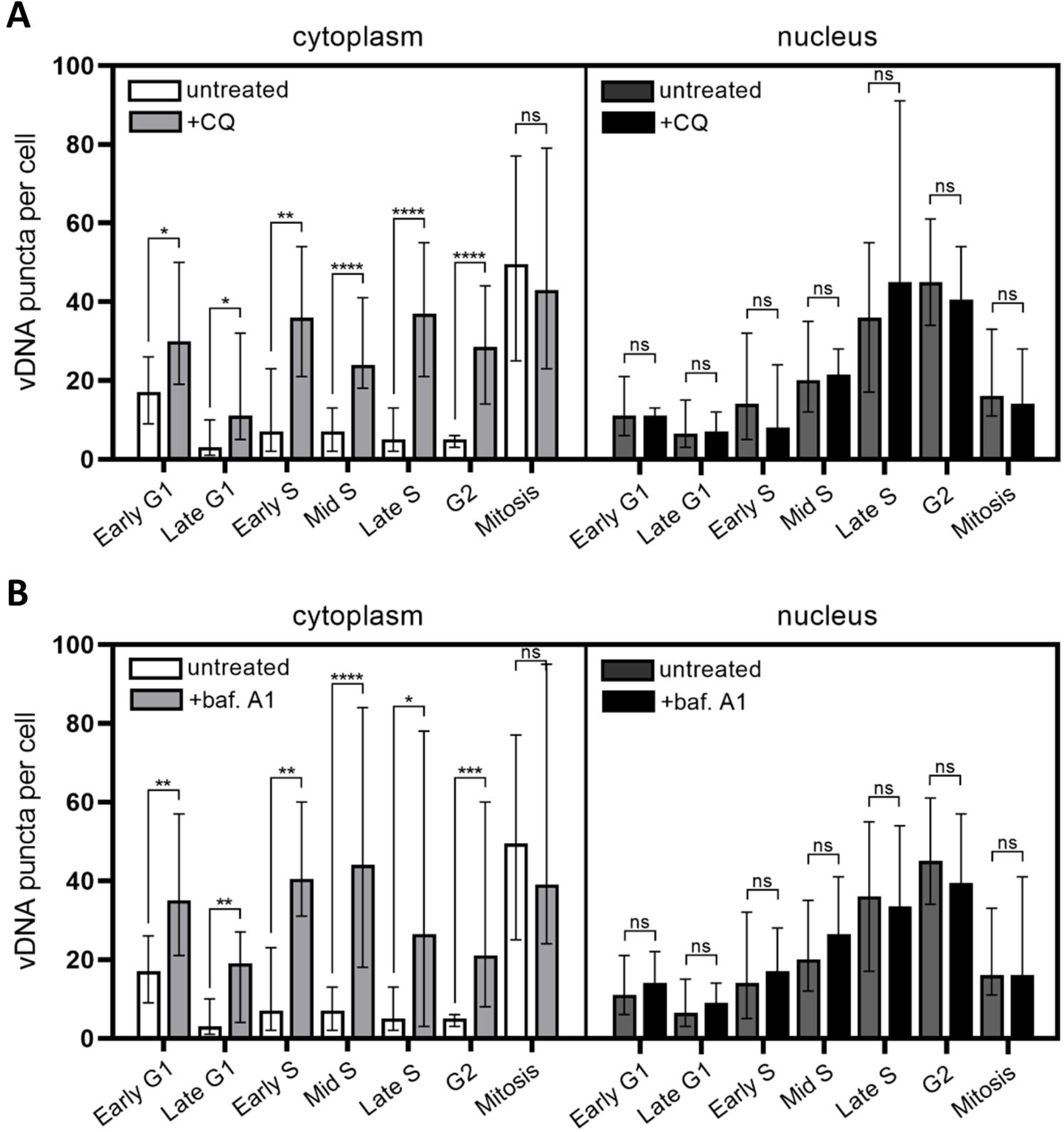
Inhibitors of lysosomal acidification block HPV genome degradation. **(A, B)** Quantification of cytoplasmic and nuclear HPV16 genome puncta in long-term (>6 weeks) infected primary foreskin keratinocytes treated with 100 μM chloroquine-(A) and 100 nM bafilomycin A1 (B). Data presented as median viral puncta with 95% CI. At least 20 cells per cell cycle stage coming from one biological experiment were analyzed.

## DISCUSSION

Herein we have provided strong evidence that viral genome maintenance in keratinocytes harboring episomal HPV is likely achieved by a balance between unlicensed genome replication (amplification) during S and G2 phase and loss of viral genome to the cytosol during mitosis; excess cytoplasmic HPV genomes are then degraded via a lysosome-dependent manner during G1 and S phase. During S and G2 phase, we observe a continuous increase in the number and size of nuclear fluorescent foci after probing for HPV genomes. This observation was made with HPV16 and HPV31 harboring cell lines independent of the time post infection, the mode of transduction, and the means of selection for outgrowth of HPV positive cell lines. Surprisingly, the majority of viral genomes fails to tether to condensed chromosomes during mitosis and ends up in the cytosol. The observed loss of viral genome to the cytosol during mitosis resets nuclear genome copy numbers to pre-S levels. Viral genome residing in the cytosol after mitosis is lost over time, suggesting it is degraded. Indeed, treatment with agents interfering with acidification of lysosomes prevents viral genome loss resulting in the accumulation of cytosolic viral genome. Overall, rather than viral genome copy number being maintained by licensed replication and being doubled in S phase to be equally distributed to the daughter cells, it cycles throughout the cell cycle (Fig 10).

**FIG 10.**
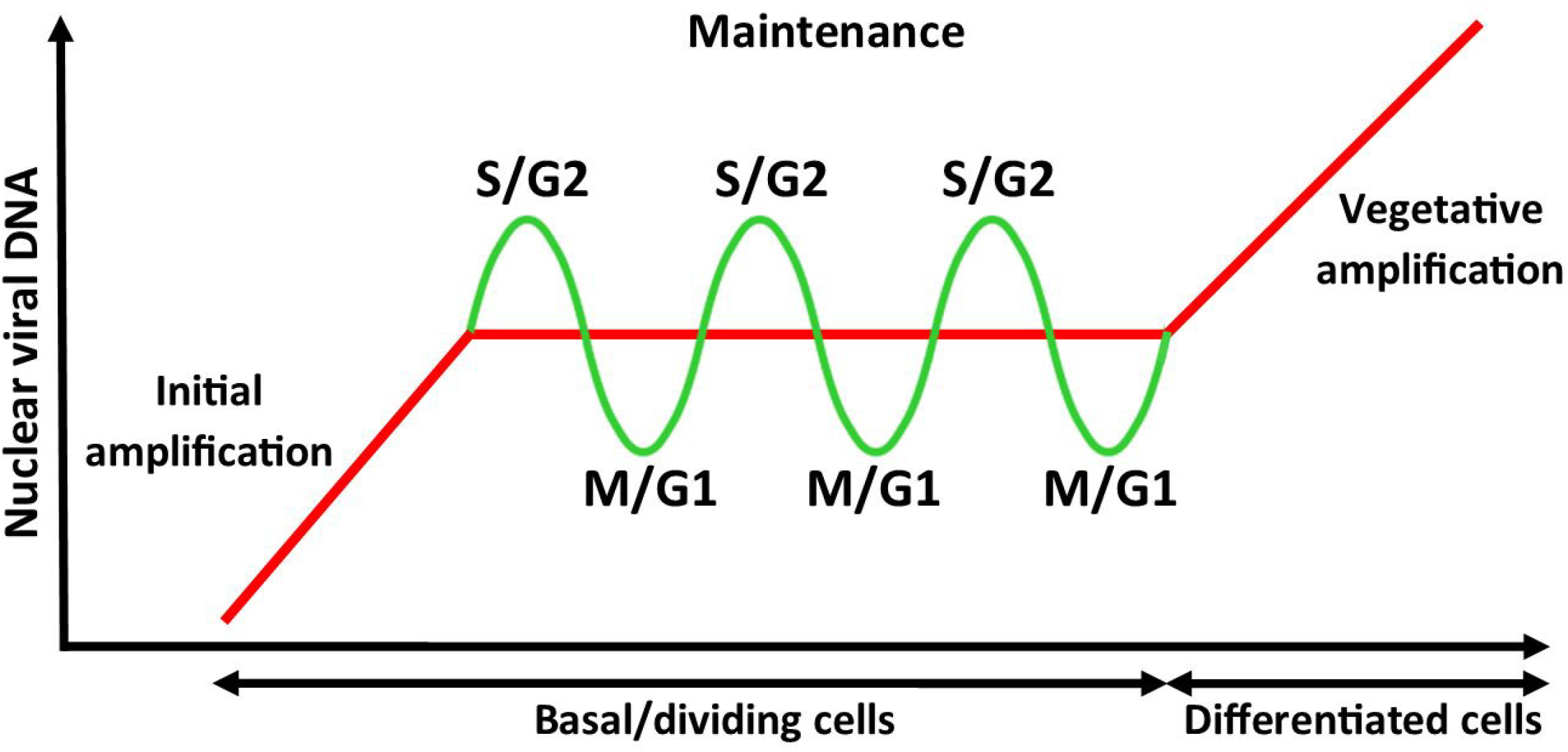
Model of papillomavirus genome amplification throughout the cell cycle. After initial amplification, genome copy number is maintained by a balance between unlicensed viral DNA amplification in S and G2 phase and subsequent loss of untethered genomes to the cytoplasm during mitosis. Cytoplasmic viral DNA is then gradually degraded over G1 and S-phase by lysosome-involving mechanism. Model modified from McBride (2).

These findings have several implications regarding our understanding of replication events during the viral lifecycle. In the accompanying manuscript by Zwolinska *et al* (), we provide strong evidence that viral genome is indeed amplified during the establishment phase in an E1 and E2-dependent manner. But rather than switching to maintenance replication, i.e., replicating viral genome once in each S phase following the establishment, the mode of replication does not change in subsequent cell cycles - HPV genome continues to be amplified. So far, no clear evidence in support of licensed DNA replication has been published. Therefore, other mechanisms have been proposed: tight control of E1 and E2 protein levels – both at the transcriptional and post-transcriptional (splicing) level (1, 3, 4, 15, 16); post-translational modification of E1 and E2 protein such as phosphorylation and proteolytic processing (3, 17, 18); and viral repressors such as the E8^E2 protein, which has been convincingly demonstrated to restrict genome copy number (5, 6, 19). Our data suggest that, while these mechanisms may be involved in genome copy number control, they are not sufficient. Instead, we propose that limited tethering to the host genome resets nuclear genome copy numbers during each mitosis. On average, we observe approximately five foci in close vicinity to mitotic chromosomes during cytokinesis. We must assume that some of genomes are randomly present without being tethered to the host cell genome. Whether restrictions to efficient tethering are due to limited viral and cellular factors or requires attachment to (a) specific site(s) on the human genome needs further studies. Putative candidates for viral and cellular tethering factors are the viral E2 protein and the host Brd4 and TopBP1 (4, 20-22). In addition, the size and intensity of the fluorescent foci suggests that each of the tethered foci contains more than one viral genome implying they are segregating in clusters.

We also observe cytoplasmic viral genome in mitotic and interphase cells in organotypic raft cultures, suggesting that limitations to viral genome tethering is not restricted to cells grown in monolayer culture. While analyses of tissue sections derived from organotypic rafts are more challenging due to their three-dimensional nature, we propose that the cycling of viral genome copy number throughout the cell cycle also occurs in tissues. This makes us wonder if we have to rethink differentiation-induced genome amplification. It has been demonstrated that HPV infected cells in the upper layers of the stratified epithelia remain in late S or G2 phase for an extended time (23, 24). In addition, E2 protein levels have been shown to increase throughout differentiation (25, 26). This likely applies also to E1 protein levels as splice variants derived from the late promoter have coding potential for E1. Others (14, 27) and we, herein, have shown that viral genome replication continuous into G2 phase. Therefore, the increased E1 and E2 protein levels in addition to extended time HPV harboring differentiated cells spend in late S and/or G2 would likely be sufficient to account for the observed differentiation induced genome amplification. While not yet experimentally proven, we find this an attractive interpretation of our data. The proposed rolling circle amplification is mainly based on two-dimensional gel analysis and enzymatic activities necessary for site-specific cutting and relegation have not been identified and are clearly not encoded by HPV viral proteins.

Cytoplasmic DNA usually triggers innate immune signaling via the cGAS/STING pathway (28). However, this is not observed in HPV harboring cells, suggesting that viral factors may interfere with this process (29). Indeed, both E6 and E7 have been shown to blunt innate immune signaling (30, 31). Alternatively, cytosolic HPV genome could be associated with histones and/or other factors that prevent cGAS binding or cGAS-mediated generation of cGAMP. However, further studies are required to investigate their role in preventing the observed cytoplasmic DNA to trigger such a response. The availability of the infection model will allow us to knock out E6 and E7 individually or together and test for cGAS/STING pathway activation under these circumstances. Further studies are also required into the mechanism by which the cytosolic DNA is degraded. Since our data suggest that lysosomal acidification is required for efficient degradation, micro-autophagy is a likely mechanism involved.

## MATERIALS AND METHODS

### Cells

HEK 293TT cells were obtained from John Schiller (NIH, Bethesda, MA) and were cultured in DMEM with non-essential amino acids, antibiotics and GlutaMAX (Gibco). Human keratinocytes HaCaT cells were purchased from the American Type Culture Collection (ATCC) and grown in low glucose Dulbecco Modified Eagle Medium (DMEM) containing 5% FBS, antibiotics, and 2.5 μg/ml of plasmocin (InvivoGen). UMSCC47 cells (Millipore Sigma, SCC071) were maintained in DMEM with GlutaMAX (Gibco, 1 g/L D-glucose, 110 mg/L sodium pyruvate) supplemented with 10% FBS (32). UMSCC47 is a head and neck squamous carcinoma cell line containing around 15-18 copies of type II integrated HPV16 (13, 32). Human neonatal primary foreskin keratinocytes (HFK) were purchased from ATCC (PSC-200-010). Three different donor derived HFKs were used in our study. HFKs were cultured in E media containing 5% FBS and 5 ng/ml of mouse epidermal growth factor (EGF, BD Biosciences; 354010) and 10 μM Rho kinase (ROCK) inhibitor Y-27632 which is known as a factor increasing HFKs lifespan (33, 34). In all experiments involving HPV16 infection, ROCK inhibitor was used up to 7 dpi, and then excluded from media. Growth of HFKs was supported by co-culture with mitomycin C-treated mouse NIH 3T3 J2 fibroblasts, as described previously (35). HPV16-WT- and HPV31-WT-positive cells established by genome transfection, as well as a HPV31-positive CIN12-9E cell line derived from a low-grade lesion were a kind gift from Jason Bodily (LSUHS, Shreveport, LA). These cells were cultured in E media containing 5% FBS and 5 ng/ml of mouse epidermal growth factor (EGF, BD Biosciences; 354010) and with NIH 3T3 J2 fibroblast added as feeders. Before harvesting cells, J2 cells were removed by short trypsin treatment and PBS wash.

### Antibodies

Antibodies used in current study were as follows: rabbit anti-cyclin B1 antibody (ab32053, Abcam), rabbit anti-MCM7/PRL (ab52489, Abcam), mouse anti-α-tubulin (3873, Cell Signaling), goat anti-rabbit-AF647 labeled secondary antibody (A21245, Invitrogen), goat anti-mouse-AF546 labeled secondary antibody (A11030, Invitrogen).

### Generation of quasiviruses

HPV16 quasivirions were generated using HEK 293TT cells following the improved protocol of Buck and Thompson (36) with minor modifications. 293TT cells were first co-transfected with the pSheLL16 L1/L2 and pEGFP-N1-HPV16 plasmids and then transfected with the pBSCre plasmid 24 hours after initial transfection. The plasmid pEGFP-N1 containing the entire floxed HPV16 genome (pEGFP-N1-HPV16) and pBSCre plasmid have been described previously (37). After two days of culture, cells were harvested and viral particles were purified as described previously (36, 38). Free DNA was removed before purification by treatment with Benzonase endonuclease (Millipore Sigma). Isolated viral particles are a mixture of pseudoviruses, serving as an internal non-amplifying control (pEGFP-N1) and quasivirions (HPV16 genomes). Such a mixture is a result of Cre recombinase activity allowing generation of two circular plasmids (pEGFP-N1 and HPV16 genome), which are subsequently packed into capsids. Viral genome equivalents (vge) were determined by qPCR of DNA isolated from encapsidated virions isolated with NucleoSpin Blood kit QuickPure (Macherey-Nagel; 740569.250).

### HPV16 infection using ECM-to-cell transfer

Infection was done as previously described (39, 40) Zwolinska 2022, accompanying paper). Briefly, extracellular matrix (ECM) was produced by HaCaT cells by seeding them on 60 mm dishes and growing them for 24-48 h until they reached confluency. HaCaT cells were then detached by 1-2 h treatment with 0.5 mM EDTA in PBS and washed out with PBS. Dish surface was treated with 8 μg/ml mitomycin C for 4 h and washed with PBS to prevent outgrowth of any residual HaCaT cells. Optiprep-purified viral particles (1-2 × 10^8^ vge/dish) diluted in 1.5 ml E medium, were then added to the ECM and incubated for 2-3 h at 37°C. Next, 1-2×10^5^ low passage HFKs were added in E media containing ROCK inhibitor. After at least 6 h post seeding of HFKs, 1×10^5^ mitomycin C-treated J2 fibroblast feeders were added. Infection was continued until HFK reached 70-80% confluence. Cells were passaged every 7-14 days depending on the growth rate and starting from 7 dpi media without ROCK inhibitor was used.

### Organotypic raft culture

3D cultures generated from HFKs infected for 5-7 days with HPV16-WT quasivirions were grown as described previously (35, 41). Collagen gel containing fibroblasts J2 feeders was used as a framework for seeding of 1×10^6^ keratinocytes. HFKs attached to the collagen gel were lifted and placed onto stainless steel grid in the culture dish. Media level was high enough to contact with the gel from below without disturbing HFKs contact with the air from above. Media was changed every other day. Rafts were grown for 21 days, and samples were then collected for *in situ* DNA hybridization and staining. Raft tissues were fixed for 30 min in 4% paraformaldehyde at 4°C, washed three times in cold PBS and once in cold 70% ethanol, and then stored in fresh cold 70% ethanol at 4°C until processing and paraffin embedding. Rafts generated from uninfected HFKs seeded on ECM were used as a control.

### Determination of cell cycle phase

Cells grown in monolayer were assigned to a certain stage of cell cycle based on 5-ethynyl-2’-deoxyuridine (EdU) incorporation (S-phase) and cyclin B1 staining. Cells negative for both EdU and cyclin B1 were recognized as G1. Distinguishing between early and late G1 was done based on the number and the size of nucleoli. Early G1 cells had small, numerous nucleoli and cells in late G1 had up to three big nucleoli. Cells in early S phase were EdU-positive and cyclin B1-negative. Mid S and late S cells were characterized by the presence of both, EdU and cyclin B1; however, EdU covered less than 50% of the nucleus of late S and more than 50% in mid S cells. G2 cells were negative for EdU and positive for cyclin B1. Mitotic cells were selected based on mitotic figures. HPV-positive and control cells growing on cover slides in monolayer culture were pulse labeled with 10 μM EdU for 30 min at 37°C. Next, they were washed twice with PBS, fixed with 4% paraformaldehyde (PFA), washed again with PBS and dehydrated with 50%, 70% and 100% ethanol gradient for 5 min each at room temperature. Slides were then rehydrated with 70% and 50% ethanol gradient for 2 min each and incubated twice with PBS for 10 min and subjected to RNA or DNAscope (see below). EdU detection was performed using Click-iT EdU Alexa Fluor 555 Imaging Kit (C10338, Invitrogen) according to the manufacturer protocol. Immunofluorescence staining was proceeded by blocking with 5% normal goat serum (NGS) for 45 min at room temperature. Slides were then incubated with cyclin B1 antibody (cat. no. ab32053, Abcam) for one hour at 37°C in humidified chamber. After extensive washing with PBS, they were incubated with AF-labeled secondary antibody in 2.5% NGS for 30 min at 37°C in humidified chamber. After another round of extensive washing with PBS slides were covered with Hoechst33342 (cat. no. C10340, Invitrogen, 1:2000 in PBS), incubated for 10 min and washed with PBS three times. After the final washing, slides were air-dried and mounted with ProLong Glass Antifade Mountant (cat. no. P36984, Invitrogen).

### Detection of viral RNA in HFKs grown in non-differentiating conditions

Viral RNA was detected using RNAscope Multiplex Fluorescent v2 kit (catalog number: 323100; Advanced Cell Diagnostics [ACD]) and a probe targeting HPV16/18 E6 and E7 (cat. no. 311121; ACD) or HPV31 (cat. no. 311551, ACD) according to the manufacturer’s protocol. Rehydrated cells on the slides were covered with pretreatment reagents (cat. no. 322381; ACD) such as hydrogen peroxide to block endogenous peroxidases and protease III at room temperature for 10 min each. After a PBS wash, samples were incubated with pre-warmed target probes for 2 h at 40°C. The positive control probe for cellular RNA detection (PPIB, cat no. 313901; ACD) provided by the supplier was also used (Fig 2A). Signal amplification and detection reagents (cat. no. 323110; ACD) were applied sequentially and incubated at 40°C in AMP1, AMP2, AMP3, and HRPC1 reagents for 30, 30, 15, and 15 min, respectively. Before adding each reagent, samples were washed twice with washing buffer (cat. no. 310091; ACD). For fluorescence detection, Alexa Fluor 488 tyramide (cat. no. T20912; Invitrogen) in RNAscope Multiplex TSA buffer (cat. no. 322809; ACD) was added for 30 min at 40°C. After PBS washing slides were covered with Hoechst33342 (1:2000 in PBS), incubated for 10 min and washed with PBS three times. After the final washing slides were air-dried and mounted with ProLong Glass Antifade Mountant. Confocal images were acquired as z-stacks (at least 12 slices per image) with the Leica TCS SP5 spectral confocal microscope using a 63x objective or Olympus CSU W1 Spinning Disk Confocal System using a 60x objective. Images were processed using Leica Application Suite X software, cellSens Software V4.1 and Imaris 9.8.2 Software.

### Detection of HPV DNA in HFKs grown in non-differentiating conditions

Detection of viral DNA was performed by using a Multiplex Fluorescent v2 kit (cat. no. 323100; ACD) and the probe targeting HPV16/18 L1 (cat no. 315601; ACD) or HPV31 (cat. no. 311551, ACD). We modified the RNAscope procedure as described by Deleage et al. (42) by adding a RNase A pretreatment step followed by short denaturation. Cells were cultured on coverslips and were fixed with 4% paraformaldehyde (PFA) for 15 min at room temperature. PFA was removed, slides were washed with PBS and then dehydrated, rehydrated, and washed with PBS again as described above. Next, slides were incubated with pretreatment reagents, as before: hydrogen peroxide and protease III at room temperature for 10 min each. Then, they were rinsed with MilliQ water and incubated with RNase A (25 μg/ml, cat no. EN0531; Thermo Scientific) for 30 min at 37°C. The RNase tissue pretreatment step was followed by a short denaturation in which the slides were incubated at 60°C with pre-warmed probes for 15 min and then transferred to 40°C for hybridization overnight. Amplification and detection were performed according to the manufacturer’s protocol, using RNAscope 0.5× washing buffer for all washing steps. After the last washing, slides were subjected to EdU detection kit and immunofluorescence staining protocol. Following PBS washing, the slides were covered with Hoechst33342 (1:2000 in PBS), incubated for 10 min and washed with PBS three times. After the final washing, slides were air-dried and mounted with ProLong Glass Antifade Mountant. Positive control probe for cellular RNA detection (PPIB, cat no. 313901; ACD), together with or without RNase A (cat. no. E866, VWR) treatment, was used for testing DNA detection specificity (Fig 2B). DNase I (cat. no. M0303L, NEB Biolabs) treatment for 30 min at 37°C was used as control of specific detection of viral DNA in described DNAscope protocol (Fig 2C). Confocal images were acquired with the Leica TCS SP5 spectral confocal microscope using a 63x objective or Olympus CSU W1 Spinning Disk Confocal System using a 60x objective. Images were processed using Leica Application Suite X software and cellSens Software V4.1. Quantifications of viral DNA puncta depending on cellular localization as well as the cell cycle phase was performed in Imaris 9.8.2 Software.

### Detection of HPV16 DNA in organotypic raft cultures by DNAscope

Viral DNA was detected with Multiplex Fluorescent v2 kit (cat. no. 323100; ACD) and the probe targeting HPV16/18 L1 (cat no. 315601; ACD). Paraffin-embedded sections of raft tissues were heated at 60°C for 1 h, deparaffinized in xylene for 10 min, followed by dehydration in 100% ethanol for 5 min before air-drying. They were then incubated with hydrogen peroxide for 10 min at room temperature. Heat-induced epitope retrieval was performed by boiling sections in RNAscope Target Retrieval Reagent for 15 min, and the sections were immediately washed with MilliQ water and then dehydrated with 100% ethanol for 5 min before air drying. Next, tissues on the slides were subjected to RNAscope Protease Plus for 30 min at 40°C. Slides were rinsed in Milli-Q water and incubated with RNase A (25 μg/ml, cat no. EN0531; Thermo Scientific) for 30 min at 37°C. The RNase tissue pretreatment step was followed by a short denaturation in which we incubated the slides at 60°C with pre-warmed probes for 15 min and then transferred them to 40°C for hybridization overnight. Amplification and detection were performed according to the manufacturer’s protocol, using RNAscope 0.5× wash buffer for all washing steps. After completing of the last washing step, tissues on the slides were covered with 2.5% NGS and stained with rabbit anti-MCM7/PRL (ab52489, Abcam) for one hour at 37°C. They were then extensively washed and incubated with anti-rabbit AF647 secondary antibodies for 30 min at 37°C. Another round of extensive washings was followed by incubation with Hoechst33342 (1:2000 in PBS) for 10 min and washed with PBS three times. After the final washing slides were air-dried and mounted with ProLong Glass Antifade Mountant. Confocal images were acquired with the Leica TCS SP5 spectral confocal microscope using a 63x objective Images were processed using Leica Application Suite X software. Quantifications of viral DNA puncta depending on cellular localization as well as the cell cycle phase, was performed using Imaris 9.8.2 Software.

### Lysosomal acidification inhibitor treatments

Infected cells grown on cover slides were incubated for 16-18 hours with the following inhibitors of lysosome acidification: 20mM NH_4_Cl, 100μM chloroquine diphosphate salt (C6628, Millipore Sigma), and 100 nM bafilomycin A1 (1334, Tocris). After treatment, slides were washed twice with PBS and fixed with 4% PFA, dehydrated with ethanol gradient and then used for DNAscope, EdU detection and immunostaining procedure as described above. Confocal images were acquired with the Leica TCS SP5 spectral confocal microscope using a 63x objective or Olympus CSU W1 Spinning Disk Confocal System using a 60x objective. Images were processed using Leica Application Suite X software and cellSens Software V4.1. Quantifications of viral DNA puncta, depending on cellular localization as well as the cell cycle phase, was performed in Imaris 9.8.2 Software.

